# HSP90α is specifically required for rod photoreceptor function and cannot be replaced by HSP90β

**DOI:** 10.1101/2025.06.05.657739

**Authors:** Juri Hoda, Hunter Aliff, Wentao Deng, Visvanathan Ramamurthy

**Author notes:** **Address for correspondence:** Visvanathan Ramamurthy, Department of Biochemistry and Molecular Medicine, West Virginia University School of Medicine; 64 Medical Center Dr. Morgantown, WV, USA, 26506.

## Abstract

Heat Shock Protein 90 (HSP90) is a critical molecular chaperone that exists as two cytosolic paralogs, HSP90α and HSP90β, which share high sequence identity but may perform non-redundant functions *in vivo*. Loss of HSP90α in mice results in progressive rod photoreceptor degeneration despite normal retinal development and expression of HSP90β. To investigate whether HSP90β can substitute for HSP90α in photoreceptors, we generated adeno-associated virus (AAV) vectors expressing HA-tagged HSP90α or HSP90β under the control of a short rhodopsin promoter. In *Hsp90α* ^*-/-*^ mice, subretinal delivery of AAV-*Hsp90aa1* (HSP90α) restored rod function and prevented photoreceptor degeneration, as measured by electroretinography (ERG). In contrast, AAV-mediated expression of HSP90β failed to rescue rod function despite comparable expression levels. Overexpression of either paralog in wild-type mice had no adverse effects on retinal function. These findings reveal a paralog-specific and intrinsic requirement for HSP90α in rod photoreceptors, demonstrating that HSP90β cannot compensate for its loss despite structural similarity.

## Introduction

Heat Shock Protein 90 (HSP90) is a highly conserved molecular chaperone critical for maintaining cellular protein homeostasis. In mammals, four HSP90 paralogs are expressed: two cytosolic paralogs (HSP90α and HSP90β), the mitochondrial paralog TRAP1, and the endoplasmic reticulum-resident GRP94. Among these, cytosolic HSP90α and HSP90β share 86% sequence identity and are often assumed to function redundantly based on *in vitro* studies, which show no clear client preference between the paralogs ^1-3^.

However, *in vivo* studies have revealed distinct, non-overlapping roles. HSP90α knockout mice exhibit male infertility despite continued expression of HSP90β, which fails to compensate for the loss of HSP90α ^1,4-7^. Conversely, HSP90β knockout results in embryonic lethality, even in the presence of HSP90α ^8^. These findings challenge the assumption of functional redundancy and raise important questions about paralog-specific roles in client protein maturation.

Indeed, several clients appear to rely specifically on one cytosolic paralog. For instance, the maturation and trafficking of the human Ether–à–go-go–Related Gene product (hERG) potassium channel depend exclusively on HSP90α ^9^, while HSP90β is required for the stability of cellular Inhibitor of Apoptosis Protein 1 (c-IAP1)^10^, and the membrane localization of specific pathogenic KCNQ4 variants ^11^. The molecular basis for this selectivity remains unclear, but emerging evidence suggests distinct client specificities and biological functions for each HSP90 paralog, with implications for targeted therapies.

In the nervous system, photoreceptor neurons of the retina provide a valuable model for investigating chaperone function. Rod and cone photoreceptors are responsible for vision in dim and bright light, respectively. Clinical trials of pan-HSP90 inhibitors in cancer patients have reported night blindness as a side effect, suggesting a role for HSP90 in rod function ^12-15^. Consistently, pharmacological inhibition of HSP90 in animal models leads to decreased levels of rod-specific proteins, including phosphodiesterase 6 (PDE6) and G-protein receptor kinase 1 (GRK1)^16^. Genetic studies in dogs and mice further support this role. We and others have shown that mice lacking HSP90α display normal retinal development and early rod function but undergo progressive rod degeneration, resulting in functional decline ^5^. Cone function, however, remains preserved. Notably, rod degeneration occurs independently of light exposure, indicating an intrinsic requirement for HSP90α for vision ^5^.

This unique postnatal dependence on HSP90α in rod photoreceptors presents an opportunity to test the functional interchangeability of cytosolic HSP90 paralogs *in vivo*. Given their high sequence similarity, we asked whether HSP90β can compensate for the loss of HSP90α in rod photoreceptors. Using adeno-associated virus (AAV)-mediated expression of HA-tagged HSP90α or HSP90β under the rod-specific rhodopsin promoter, we evaluated their ability to restore rod function in *Hsp90α*-deficient mice. Our findings reveal that while HSP90α expression preserves photoreceptor function, equivalent levels of HSP90β do not. These results demonstrate a unique and non-redundant requirement for HSP90α in the maintenance of rod photoreceptors.

## Results

### Photoreceptor function is unchanged by AAV-driven HSP90 overexpression

A whole-body knockout of *Hsp90aa1* (HSP90α) exhibited two major phenotypes: male infertility and vision loss. Interestingly, retinae lacking HSP90α developed normally and initially displayed normal photoreceptor function, as measured by electroretinography (ERG). However, rod photoreceptor neurons, progressively undergo degeneration, leading to complete loss by six months of age. In contrast, cone photoreceptor cells do not show a dependency on HSP90α. These findings raised an important question: Is HSP90α intrinsically required in rod photoreceptor cells, and if so, can its close paralog *Hsp90ab1* (HSP90β) functionally substitute for the loss of HSP90α?

To address these questions, we generated Adeno-Associated Virus (AAV) vectors using a Y733F mutant of AAV serotype 8 that expresses either mouse HSP90α (*Hsp90aa1*) or HSP90β (*Hsp90ab1*). We used AAV8 (Y733F) because this serotype efficiently transduces photoreceptor cells ^17,18^. In addition, we expressed the transgene under a short rod opsin promoter (Rhop). To ensure comparable expression levels, both constructs were tagged at their N-terminus with the Haemagglutinin (HA) epitope tag (Fig. 1A).

**Figure 1.**
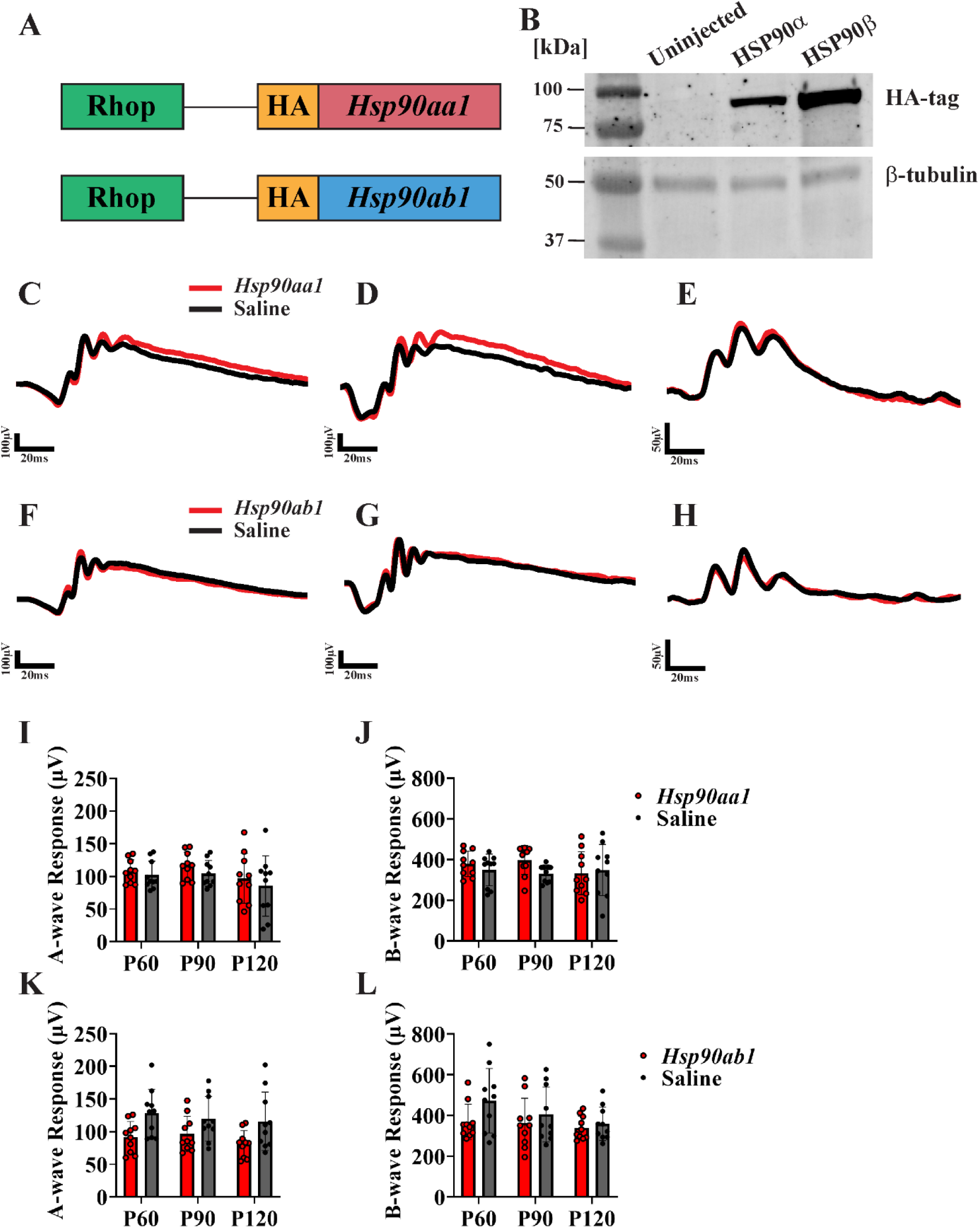
Expression of Hsp90 in rod photoreceptors does not alter visual function in wild-type mice. **A**. Schematic representation of the AAV-*Hsp90aa1* and AAV-*Hsp90ab1* constructs. Each vector contains a rhodopsin promoter driving expression of HA-tagged HSP90α or HSP90β. **B**. Immunoblot analysis of retinal lysates from wild-type mice at P70, uninjected or injected with AAV-*Hsp90aa1* or AAV-*Hsp90ab1*. Blots were probed with antibodies against the HA-tag and β-tubulin. Molecular weight markers (kDa) are shown on the left. **C-D**. Representative scotopic traces at P120 from wild-type mice injected with AAV-*Hsp90aa1* (red) or saline (black), recorded at light intensities of 0.158 cd·s/m^2^ (**C**) and 2.5 cd·s/m^2^ (**D**). **E**. Representative photopic trace at P120 from wild-type mice injected with AAV-*Hsp90aa1* (red) or saline (black), measured at 25 cd·s/m^2^. **F-G**. Representative scotopic ERG traces at P120 from wild-type mice injected with AAV-*Hsp90ab1* (red) or saline (black), recorded at 0.158 cd·s/m^2^ (**F**) and 2.5 cd·s/m^2^ (**G**). **H**. Representative photopic trace at P120 from wild-type mice injected with AAV-*Hsp90ab1* (red) or saline (black), measured at 25 cd·s/m^2^. **I-J**. Quantification of scotopic A-wave (**I**) and B-wave (**J**) amplitudes at P60, P90, and P120 from wild-type mice injected with AAV-*Hsp90aa1* (red) or saline (black). **K-L**. Quantification of scotopic A-wave (**K**) and B-wave (**L**) amplitudes from wild-type mice injected with AAV-*Hsp90ab1* (red) or saline (black). ERG recordings were obtained from 10 mice per group, including both male and female animals. Statistical comparisons were performed using two-way ANOVA with Tukey post hoc test (GraphPad Prism). Unlabeled comparisons were not statistically significant.

First, we tested if AAV transgenesis in wild-type mice led to the expression of HSP90. After the subretinal injection of AAV on postnatal day 21 (P21), the retinal tissue was harvested at P70. Immunoblotting of retinal lysates with an antibody against the HA antibody showed robust expression of HSP90α and HSP90β in injected eyes. In contrast, the uninjected eye or saline-injected eyes showed no expression of HA-tagged protein. β-tubulin levels served as a loading control (Fig. 1B).

Next, we tested whether the overexpression of HSP90 affects visual function using ERG at post-injection on days P60, P90, and P120. In ERG recordings, the A-wave reflects photoreceptor activity, while the B-wave corresponds to responses from downstream ON-bipolar cells. At low scotopic ranges, reflecting rod responses, and at higher intensities, representing mixed rod-cone responses, overexpression of HSP90α did not alter ERG responses compared to saline-injected eyes (At P120, A_mean_ = 97.19 µV – HSP90α, A_mean_ = 85.42 µV – Saline, n=10, *p* = 0.9462) (Fig1C-D). Similarly, the photopic or cone responses remained unaffected (Fig. 1E). Both A- and B-wave amplitudes remained stable through P120 (4 months) (Fig. 1 I-J).

When we investigated the impact of HSP90β overexpression, we found, as with HSP90α, that scotopic and photopic ERG responses showed no significant changes compared to saline controls (Fig. 1F-H). While a mild reduction in scotopic responses was observed, the difference was not statistically significant (At P120, A_mean_ = 81.26 µV – HSP90β, A_mean_ = 115.8 µV – Saline, n=10, *p* = 0.1714) (Fig. 1K-L).

Together, these findings demonstrate that AAV-mediated overexpression of HSP90α or HSP90β in rod photoreceptors is robust and non-toxic. Overexpression of either cytosolic HSP90 paralogs does not impair retinal function, supporting the utility of this system for testing functional redundancy *in vivo*.

### HSP90α expression in *Hsp90α*^*-/-*^photoreceptors restores visual responses

Next, we investigated whether the expression of HSP90α in the photoreceptors of HSP90α^-/-^ mice could restore function. AAV vectors expressing HA-tagged HSP90α (*Hsp90aa1*) were injected into one eye at P21, while the contralateral eye received an equal volume of saline as a control.

When we checked the expression of exogenous HSP90α by AAV transduction, immunoblot analysis at P70 confirmed robust expression of HA-tagged HSP90α in AAV-injected eyes, with protein migrating at the expected molecular weight. In contrast, HA-signal was not detected in saline-injected or uninjected controls (Fig. 2A).

**Figure 2.**
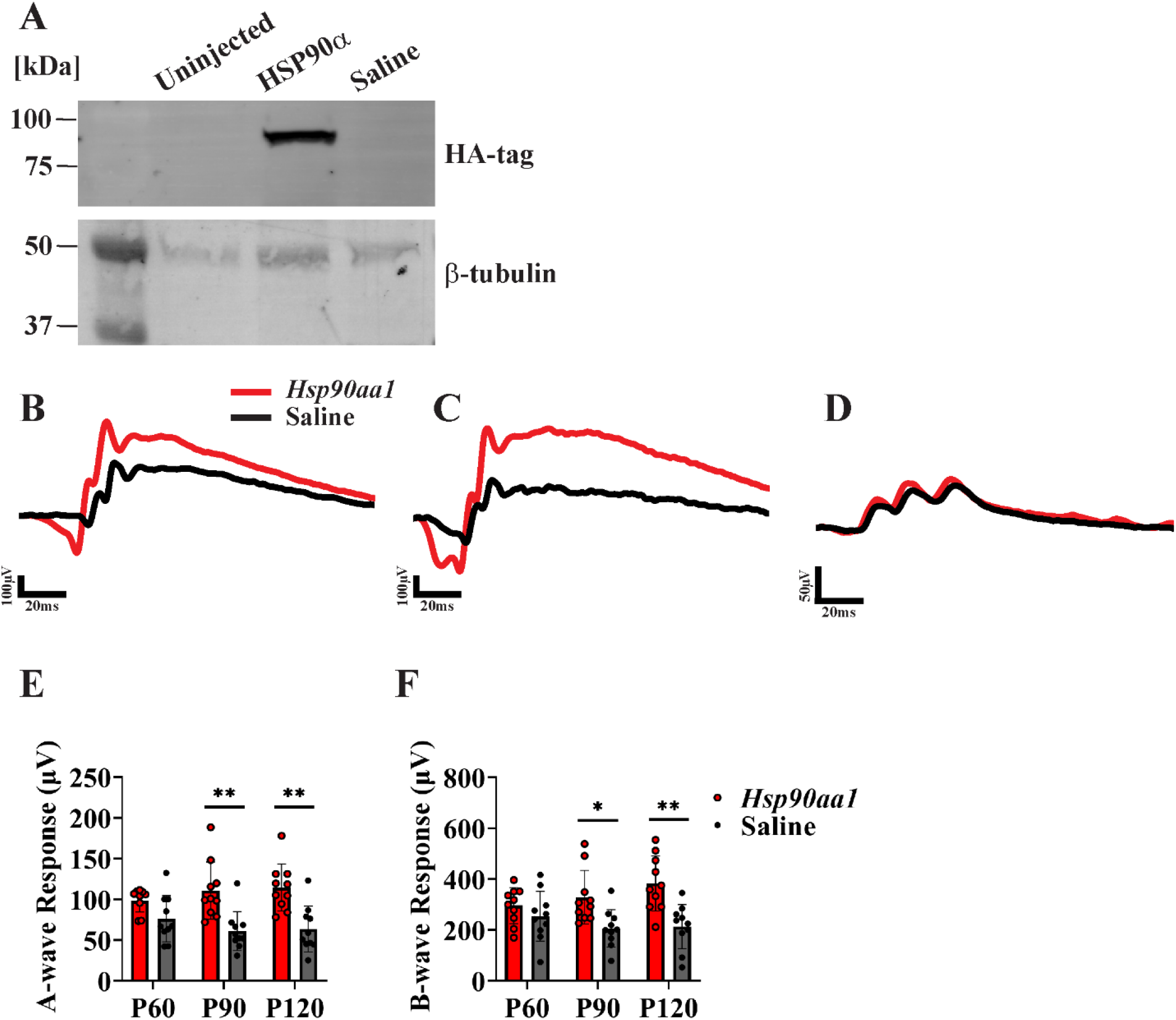
Expression of *Hsp90aa1* in rod photoreceptors restores visual function in animals lacking HSP90α. **A**. Immunoblot analysis of retinal samples from *Hsp90α*^-/-^ mice at P70, comparing uninjected, AAV-*Hsp90aa1* injected eyes, and saline-injected controls. Blots were probed with antibodies against the HA-tag and β-tubulin. Molecular weight (kDa) standards are indicated on the left. **B-C**. Representative scotopic ERG traces at P120 from *Hsp90α*^*-/-*^ treated with AAV-*Hsp90aa1* (red) or saline (black), recorded at light intensities of 0.158 cd·s/m^2^ (**B**) and 2.5 cd·s/m^2^ (**C**). **D**. Representative photopic ERG trace at P120 from *Hsp90α*^-/-^ mice injected with AAV-*Hsp90aa1* (red) or saline (black), measured at 25 cd·s/m^2^. **E-F**. Quantification of scotopic-A wave (**E**) and B-wave (**F**) amplitudes at P60, P90, and P120, comparing AAV-*Hsp90aa1* (red) treated eyes with saline-injected controls (black) in *Hsp90α*^-/-^ mice. ERG recordings were obtained from 10 mice per group, including both male and female animals. Statistical analyses were performed using two-way ANOVA with Tukey’s post hoc test in GraphPad Prism. Significance was defined as *p* ≤ 0.05 (*) and *p* ≤ 0.01 (**). Unlabeled comparisons were not statistically significant.

We then assessed whether AAV-mediated expression of HSP90α in mature photoreceptors in the context of whole-body HSP90α deficiency. ERG responses were recorded post-injection on days P60, P90, and P120. As expected, saline-injected or uninjected *Hsp90α*^*-/-*^ mice exhibited progressive loss of scotopic responses. In contrast, AAV-*Hsp90aa1* injected eyes exhibited a robust rescue of rod photoreceptor function (Fig. 2B-C). This functional improvement was sustained and continued to improve over time, whereas visual function in saline or untreated eyes declined (At P120, A_mean_ = 114.4 µV – HSP90α, A_mean_ = 63.56 µV – Saline, n=10, p = 0.0014) (Fig. 2E-F).

Consistent with our previous findings, photopic ERG responses were unaffected by the absence or restoration of HSP90α, indicating that cone function remains independent of HSP90α levels (Fig. 2D).

In summary, these results demonstrate that expression of HSP90α, specifically in photoreceptors, is sufficient to restore rod-mediated visual function in *Hsp90α*^*-/-*^ mice. This finding highlights the intrinsic requirement for HSP90α in maintaining long-term rod photoreceptor function and vision.

### HSP90β is unable to functionally compensate for HSP90α loss in rod photoreceptors

We next tested whether HSP90β could substitute for HSP90α to sustain photoreceptor function. Using a similar strategy, we subretinally administered AAV expressing HA-tagged *Hsp90ab1* in one eye of *Hsp90α*^*-/-*^ mice at postnatal day 21 (P21). The contralateral eye received saline injection as a control.

Immunoblotting of retinal lysates at P70 confirmed robust expression of HA-HSP90β at the expected molecular weight in AAV-injected eyes, while no signal was detected in saline-injected or uninjected controls (Fig. 3A).

**Figure 3.**
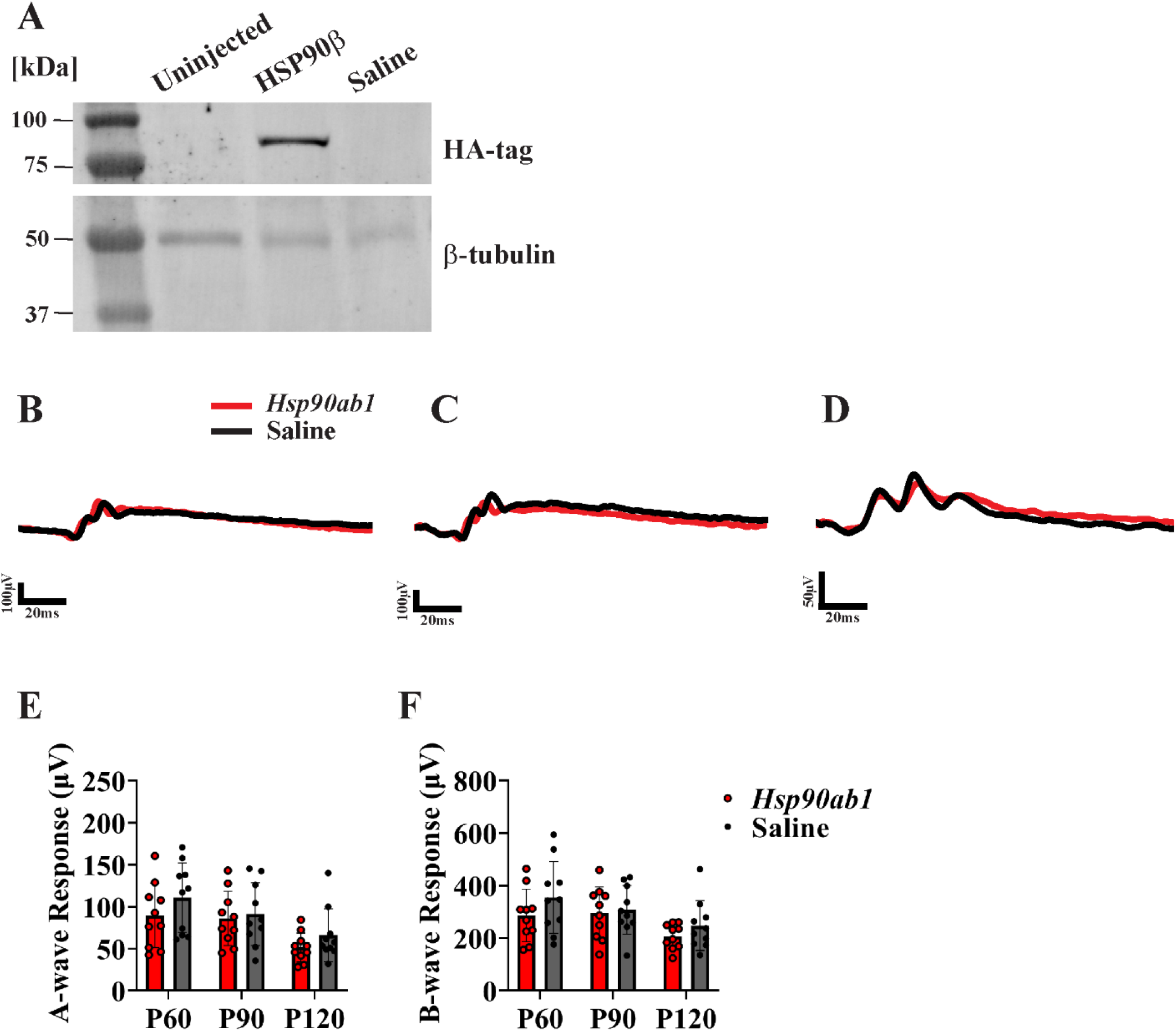
Expression of *Hsp90ab1* in rod photoreceptors fails to rescue visual function in *Hsp90α*^*-/-*^mice. **A**. Immunoblot of retinal lysates from *Hsp90α*^*-/-*^ mice at P70, including an uninjected eye, an eye treated with AAV-*Hsp90ab1*, and a saline-injected control. Blots were probed with antibodies against the HA-tag and β-tubulin. Molecular weight markers (kDa) are shown on the left. **B-C**. Representative scotopic traces at P120 from *Hsp90α*^*-/-*^ mice injected with AAV-*Hsp90ab1* (red) or saline (black), recorded at light intensities of 0.158 cd·s/m^2^ (**B**) and 2.5 cd·s/m^2^ (**C**). **D**. Representative photopic ERG trace at P120 from *Hsp90α*^*-/-*^ mice injected with AAV-*Hsp90ab1* (red) or saline (black), measured at 25 cd·s/m^2^. **E-F**. Quantitative analysis of scotopic A-wave (E) and B-wave (**F**) amplitudes plotted at P60, P90, and P120, comparing injected AAV-*Hsp90ab1* (red) to saline-injected controls (black) in *Hsp90α*^*-/-*^mice. ERG recordings were obtained from 10 mice per group, including both male and female animals. Statistical analysis was performed using two-way ANOVA with Tukey’s post hoc test (GraphPad Prism). Unlabeled comparisons were not statistically significant.

To assess functional recovery, ERG responses were recorded at P60, P90, and P120. At all time points tested, AAV-mediated expression of HSP90β failed to restore scotopic responses in *Hsp90α*^*-/-*^ mice (Fig. 3 B-C, E-F). Scotopic ERG traces from AAV-*Hsp90ab1* treated eyes closely resembled those from saline-injected controls, with both groups exhibiting similarly decreased A- and B-wave amplitudes (At P120, A_mean_ = 51.39 µV – HSP90β, A_mean_ = 65.88 µV – Saline, n=10, p = 0.93) (Fig. 3B-C). Moreover, similar to saline-injected controls, the scotopic responses over time declined in AAV-*Hsp90ab1*injected eyes, indicating progressive deterioration of rod function. The photopic responses remained unchanged between groups, as expected, given that cone function is not dependent on HSP90α (Fig. 3D).

These findings indicate that, despite expressing HSP90β at a similar levels as HSP90α in *Hsp90α* ^*-/-*^ mice, HSP90β is unable to compensate for the loss of HSP90α in rod photoreceptors. Our findings support a non-redundant, paralog-specific requirement for HSP90α in sustaining rod photoreceptor function.

## Discussion

HSP90 is a highly conserved molecular chaperone essential for the folding, stability, and function of client proteins. In mammals, the two cytosolic paralogs, HSP90α (*Hsp90aa1*) and HSP90β (*Hsp90ab1*), share ∼86% sequence identity and have long been assumed to be functionally redundant based on *in vitro* studies ^1-3^. However, *in vivo*, evidence challenges this assumption, particularly in the context of cell-type-specific demands ^1,4-7^. In this study, we demonstrate that HSP90α is intrinsically required for maintaining rod photoreceptor function. HSP90α performs a unique, non-redundant function in rod photoreceptors and cannot be substituted by its close paralog, HSP90β.

Our work, as well as that of others, has demonstrated that the global loss of HSP90α results in progressive rod degeneration despite normal photoreceptor development and unaltered early visual function. These findings indicated a post-developmental requirement for HSP90α in maintaining photoreceptors. In this work, we build on those observations by using AAV-mediated gene delivery to test the functional interchangeability of cytosolic HSP90 paralogs *in vivo*. Subretinal injection of AAV-*Hsp90aa1* in *Hsp90α*^*-/-*^ mice rescued rod function and this rescue was sustained, demonstrating that re-expression of HSP90α in mature photoreceptors is sufficient to restore visual function.

In contrast, overexpression of HSP90β in *Hsp90α*^*-/-*^ mice failed to rescue rod function despite robust expression and the use of an identical promoter and vector backbone. This finding underscores a paralog-specific requirement for HSP90α in rod photoreceptors and argues against functional redundancy between the two cytosolic paralogs in rod photoreceptor neurons. While both HSP90α and HSP90β can bind many of the same client proteins *in vitro*, several studies now support client-specific preferences *in vivo*. For example, the hERG potassium channel requires HSP90α for maturation, while c-IAP1 and mutant KCNQ4 trafficking rely on HSP90β. Our results extend these observations by identifying rod photoreceptors as a cell type that depends explicitly on HSP90α.

The mechanistic basis for this paralog specificity remains unclear. One possibility is that HSP90α may preferentially associate with rod photoreceptor-specific co-chaperones or post-translational modifications that HSP90β cannot mimic. Alternatively, HSP90α may stabilize a subset of rod-specific client proteins essential for phototransduction, such as phosphodiesterase 6 (PDE6) ^5,19^. Elucidating these client-specific interactions and the role of cochaperones will be an important direction for future studies.

Importantly, we also show that overexpression of either HSP90α or HSP90β in wild-type retinas does not perturb photoreceptor function, suggesting that both constructs are well tolerated *in vivo* and do not exert toxic effects. This observation strengthens the therapeutic relevance of our AAV-based approach and supports the potential for paralog-selective chaperone modulation in the treatment of retinal degenerative diseases.

In summary, our study provides compelling *in vivo* evidence that HSP90α is required in rod photoreceptors for long-term functional maintenance and cannot be replaced by its highly similar paralog HSP90β. These findings underscore the significance of paralog specificity in chaperone biology, providing new insights into the cellular selectivity of HSP90 paralog function in neurons.

### Animal model and genotyping

We utilized *Hsp90aa1* knockout (HSP90α^-/-^) mice, which were previously developed in our laboratory. This HSP90α^-/-^ model was generated by CRISPR/Cas9 genome editing. Guide RNAs were selected based on their predicted efficiency and minimal off-target effects (http://crispr.mit.edu/). Synthetic single-guide RNAs (sgRNAs) were synthesized using the CRISPR Evolution sgRNA EZ Kit (Synthego). Two sgRNAs, 5′-CCACAATCCTCTTCAGATACCAC-3′, and 5′-CCTGAAGCTCCCTTTAGATTAA-3′, were designed to target intronic regions flanking exon 4 of the *Hsp90aa1* gene (ENSMUSE00001252094). These sgRNAs were co-injected with Cas9 nuclease into mouse blastocysts to generate founders. Founders were screened by sequencing to identify heterozygous animals with exon 4 deletion, and these were backcrossed to a C57BL/6J background. All mice carried the hypomorphic *Rpe65* allele, which encodes methionine at amino acid position 450.

Genotyping was conducted using genomic DNA extracted from ear punch samples. Genotyping of *Hsp90aa1* was performed using a multiplex PCR approach with two primer sets. The first set, consisting of forward primer 5′-CAATTGTAGGGGGTGTCTGG-3′ and reverse primer 5′-TCCTCCTCTTCTTCATCAGAGC-3′, flanked exon 4, producing amplicons of 820 base pairs for *Hsp90α*^+/+^ and 336 base pairs for *Hsp90α*^-/-^. The second primer set, forward primer 5′-TTTGTGGGGAAGGTTAGCTG-3′ and reverse primer 5′-TGGGATAGCCAATGAACTGA-3 ′, specifically amplified the wild-type allele.

To generate *Hsp90α*^*-/-*^ mice carrying the *Rpe65* L450 allele, heterozygous HSP90α mice were backcrossed onto a 129SV/E background (Charles River Laboratories, Straine #287). This strain carries the *Rpe65* allele, which encodes leucine at residue 450 and expresses a normal level of RPE65 protein. Genotyping for the *Rpe65* allele was performed using standard PCR amplification with Quick-Load Taq DNA polymerase (New England Biolabs, Ipswich, MA, USA). To distinguish between the RPE65 methionine/methionine (M/M) and leucine/leucine (L/L) alleles, the following primers were used: forward primer 5′-TTACCAGAAATTTGGAGGGAAA-3′ and reverse primer 5′-CAGAGCATCTGGTTGAGAAACA-3′. The amplified products were subsequently submitted to Psomagen Inc. (Rockville, MD, USA) for Sanger sequencing.

The animals used in this study were raised under 12-hour light/12-hour dark cycles, with food and water provided *ad libitum*. All experimental procedures involving animals in this study were approved by the Institutional Animal Care and Use Committee at West Virginia University. Both female and male mice were used in this study.

### AAV vectors

The adeno-associated virus (AAV) vectors used in this study were engineered to express either HSP90α or HSP90β, tagged with an HA epitope, under the control of the rhodopsin promoter, enabling rod-specific expression. Vectors were packaged using the AAV 2/8-Y733F capsid and purified following established protocols at the West Virginia University Biochemistry and Molecular Medicine viral core facility.

### Subretinal injection

Subretinal injections were performed at postnatal day 21 (P21). AAV solutions were mixed with fluorescein dye (final concentration: 0.1%) and adjusted to a titer of 1 × 10^10^ vector genomes/µL. Pupils were dilated with a 1:1 solution of 8% tropicamide and 1.5% phenylephrine hydrochloride. Mice were anesthetized with ketamine (80 mg/kg) and xylazine (10 mg/kg) administered intramuscularly in sterile PBS. A 25-gauge needle was used to make a corneal entry, followed by a subretinal injection of 1 µL AAV using a 33-gauge blunt needle attached to a 5 µL Hamilton syringe. The contralateral eye received a saline injection as a control. Post-injection, Neomycin/Polymyxin B/Bacitracin ophthalmic ointment (Bausch & Lomb, Tampa, FL, USA) was applied. Anesthesia was reversed with an intraperitoneal injection of Antisedan (Orion Corporation, Espoo, Finland).

### Immunoblotting

Mice were euthanized via carbon dioxide (CO_2_) inhalation followed by cervical dislocation. Eyes were promptly enucleated, the cornea and lens were removed, and retinas were carefully dissected, immediately flash-frozen, and stored at −80°C until needed. Retinal tissue was homogenized through sonication in a lysis buffer composed of 0.1% Triton X-100, 50 mM Tris (pH 7.5), 300 mM NaCl, 5 mM EDTA, and a protease inhibitor cocktail (Roche). The lysates were centrifuged at 12,000 × g for 5 minutes at 4 °C, and the cellular debris was removed. Protein concentrations were determined using a BCA protein assay kit (Thermo Fisher Scientific). To prepare samples for SDS-PAGE, protein lysates were mixed with 1× SDS sample buffer (containing 2% SDS, 10% glycerol, 5% 2-mercaptoethanol, 0.002% bromophenol blue, and 62.5 mM Tris, pH 6.8). Then, they were boiled for 5 minutes. For each sample, 50 μg of total protein was loaded onto SDS-PAGE gels for electrophoretic separation. Proteins were transferred to polyvinylidene difluoride (PVDF) membranes (Immobilon-FL; Millipore).

After transfer, membranes were stained to visualize total protein (LI-COR Biosciences), which was also used for normalization and quantification. Membranes were then blocked with Odyssey Blocking Buffer (LI-COR Biosciences) for 30 minutes at room temperature. Primary antibodies, anti-mouse HA antibody (RRID: AB_2565335) and anti-rabbit β-tubulin (RRID: AB_2303998), were used in a 1:1 mixture of blocking buffer (Rockland) and PBS-T (PBS with 0.1% Tween-20), and the membranes were incubated overnight at 4°C on a bidirectional rotator. Following primary antibody incubation, membranes were washed three times with PBS-T, each for 5 minutes, and then incubated for 1 hour at room temperature with secondary antibodies at a 1:50,000 dilution in PBS-T, including goat anti-mouse Alexa Fluor 488 and goat anti-rabbit Alexa Fluor 680 or 488 (Invitrogen). After a final series of three 5-minute washes in PBS-T, membranes were imaged using the Amersham Typhoon (Cytiva). Densitometry was performed using FIJI/ImageJ, and expression levels were normalized to total protein and tubulin.

### Electroretinography (ERG)

ERG was conducted on mice at P60, 90, and 120. Following overnight dark adaptation, mice were anesthetized using 2.0% isoflurane delivered in oxygen at a flow rate of 2.5 L/min. Pupil dilation was performed with a 1:1 mixture of 8% tropicamide and 1.5% phenylephrine hydrochloride. During this experiment, animals were placed on a temperature-controlled platform maintained at 37 °C and received a continuous supply of isoflurane (1.5%) in oxygen via a nose cone. ERG recordings were obtained using silver wire electrodes positioned on the corneal surface of both eyes, with contact facilitated by a drop of 2% hypromellose in PBS (Gonioscopic Prism Solution; Wilson Ophthalmic, Mustang, OK, USA) to protect the cornea and ensure stable electrode contact. A reference electrode was inserted subcutaneously into the scalp between the ears.

Scotopic responses were recorded in complete darkness using white LED light flashes ranging from 2.45 × 10^−4^ to 2.5 cd·s/m^2^. After dark-adapted responses were collected, mice were exposed to a white background light (30 cd/m^2^) for 10 minutes to saturate rod photoreceptors. After light adaptation, recordings for photopic responses were collected using white light flashes in the presence of rod-saturating conditions. All measurements were acquired using the UTAS Visual Diagnostic System equipped with the BigShot Ganzfeld dome, UBA-4200 amplifier/interface, and EMWIN software (version 9.0.0; LKC Technologies, Gaithersburg, MD, USA).

## Data and statistical analysis

Quantitative comparisons were carried out between saline-injected and contralateral AAV-injected groups. Each experimental group, consisting of either HSP90α^-/-^ or HSP90α^+/+^ treated with AAV-HA-*Hsp90aa1* or AAV-HA-*Hsp90ab1*, included at least 10 animals, with both males and females represented. Statistical evaluation of the data was performed using GraphPad Prism version 7.0, and results are expressed as the mean ± standard error of the mean (SEM). Differences in ERG responses were assessed using a two-way ANOVA with the Tukey post hoc test, with statistical significance set at *p ≤ 0.05 or \*\**p* ≤ 0.01. Densitometric analysis of immunoblot data was performed using FIJI/ImageJ. Figures were generated using BioRender and Adobe Illustrator.

## CRediT authorship contribution statement

**Juri Hoda:** Conceptualization, Methodology, Investigation, Formal analysis, Data curation, Writing – original draft, Writing – review & editing. **Hunter Aliff:** Conceptualization, Investigation, Methodology, Writing –editing. **Wentao Deng:** Conceptualization, Investigation, Methodology, Writing – review & editing. **Visvanathan Ramamurthy:** Conceptualization, Methodology, Data curation, Project Administration, Funding acquisition, Writing – review & editing.

## Funding and support

This work was supported by the National Institutes of Health grants R01 EY031346 (VR), R01 EY028035 (VR), and P20 GM144230 (VR) from the National Institute of General Medical Sciences (NIGMS); an unrestricted challenge grant from Research to Prevent Blindness (RPB); the West Virginia Lions Club Foundation; and the Lions Clubs International Foundation.

## Data availability statement

Data will be made available upon request.

## Declaration of interest

The authors declare no conflicts of interest.

## Notes

### Competing Interest Statement

The authors have declared no competing interest.

